# A comprehensive reanalysis of publicly available GWAS datasets reveals an X chromosome rare regulatory variant associated with high risk for type 2 diabetes

**DOI:** 10.1101/112219

**Authors:** Sílvia Bonás-Guarch, Marta Guindo-Martínez, Irene Miguel-Escalada, Niels Grarup, David Sebastian, Elias Rodriguez-Fos, Friman Sánchez, Mercé Planas-Félix, Paula Cortes-Sánchez, Santi González, Pascal Timshel, Tune H Pers, Claire C. Morgan, Ignasi Moran, Juan R González, Ehm A. Andersson, Carlos Díaz, Rosa M. Badia, Miriam Udler, Jason Flannick, Torben Jørgensen, Allan Linneberg, Marit E. Jørgensen, Daniel R. Witte, Cramer Christensen, Ivan Brandslund, Emil V. Appel, Robert A. Scott, Jian’an Luan, Claudia Langenberg, Nicholas J. Wareham, InterAct Consortium, The SIGMA T2D consortium, Oluf Pedersen, Antonio Zorzano, Jose C Florez, Torben Hansen, Jorge Ferrer, Josep Maria Mercader, David Torrents

**Affiliations:** Barcelona Supercomputing Center (BSC). Joint BSC-CRG-IRB Research Program in Computational Biology, 08034 Barcelona, Spain.; Genomic Programming of Beta-cells Laboratory, Institut d’Investigacions August Pi i Sunyer (IDIBAPS), 08036 Barcelona, Spain.; Instituto de Salud Carlos III, Centro de Investigación Biomédica en Red de Diabetes y Enfermedades Metabólicas Asociadas (CIBERDEM), Madrid, Spain.; Department of Medicine, Imperial College London, London W12 0NN, United Kingdom; The Novo Nordisk Foundation Center for Basic Metabolic Research, Section for Metabolic Genetics, Faculty of Health and Medical Sciences, University of Copenhagen, Copenhagen, 2100, Denmark.; Institute for Research in Biomedicine (IRB Barcelona), The Barcelona Institute of Science and Technology, Baldiri Reixac, 10-12, 08028 Barcelona, Spain.; Departament de Bioquímica i Biomedicina Molecular, Facultat de Biologia, Universitat de Barcelona, 08028 Barcelona, Spain.; Computer Sciences Department, Barcelona Supercomputing Center (BSC-CNS), 08034 Barcelona, Spain.; Department of Epidemiology Research, Statens Serum Institut, Copenhagen, 2300 Denmark.; Division of Endocrinology and Center for Basic and Translational Obesity Research, Boston Children’s Hospital, Boston, 02116, USA.; Medical and Population Genetics Program, Broad Institute of MIT and Harvard, Cambridge, 0214, USA.; ISGlobal, Centre for Research in Environmental Epidemiology (CREAL).; CIBER Epidemiología y Salud Pública (CIBERESP).; Universitat Pompeu Fabra (UPF).; Artificial Intelligence Research Institute (IIIA), Spanish Council for Scientific Research (CSIC).; Programs in Metabolism and Medical & Population Genetics, Broad Institute of Harvard and MIT, Cambridge, Massachusetts, USA.; Diabetes Unit and Center for Genomic Medicine, Massachusetts General Hospital, Boston, Massachusetts, USA.; Department of Molecular Biology, Harvard Medical School, Boston, Massachusetts, USA.; Research Centre for Prevention and Health, Capital Region of Denmark, Copenhagen, Denmark.; Faculty of Health and Medical Sciences, University of Copenhagen, Copenhagen, Denmark.; Faculty of Medicine, University of Aalborg, Aalborg, Denmark.; Department of Clinical Experimental Research, Rigshospitalet, Glostrup, Denmark.; Department of Clinical Medicine, Faculty of Health and Medical Sciences, University of Copenhagen, Copenhagen, Denmark.; Steno Diabetes Center, Gentofte, Denmark.; National Institute of Public Health, Southern Denmark University, Denmark.; Department of Public Health, Aarhus University, Aarhus, Denmark.; Danish Diabetes Academy, Odense, Denmark.; Medical department, Lillebaelt Hospital, Vejle, Denmark.; Department of Clinical Biochemistry, Lillebaelt Hospital, Vejle, Denmark.; Institute of Regional Health Research, University of Southern Denmark, Odense, Denmark.; MRC Epidemiology Unit, University of Cambridge School of Clinical Medicine, Cambridge Biomedical Campus, Cambridge, CB2 0QQ, UK.; Members of the consortia are provided in Appendix S1.; Department of Medicine, Harvard Medical School, Boston, Massachusetts, USA.; Faculty of Health Sciences, University of Southern Denmark, Odense, Denmark.; Institució Catalana de Recerca i Estudis Avançats (ICREA).

**Keywords:** dbGaP, publicly available GWAS data, data sharing, genotype imputation, type 2 diabetes, rare variants, 1000 Genomes, UK10K, data sharing, enhancer, AGTR2, angiotensin receptor 2

## Abstract

The reanalysis of publicly available GWAS data represents a powerful and cost-effective opportunity to gain insights into the genetics and pathophysiology of complex diseases. We demonstrate this by gathering and reanalyzing public type 2 diabetes (T2D) GWAS data for 70,127 subjects, using an innovative imputation and association strategy based on multiple reference panels (1000G and UK10K). This approach led us replicate and fine map 50 known T2D *loci*, and identify seven novel associated regions: five driven by common variants in or near *LYPLAL1, NEUROG3, CAMKK2, ABO* and *GIP* genes; one by a low frequency variant near *EHMT2;* and one driven by a rare variant in chromosome Xq23, associated with a 2.7-fold increased risk for T2D in males, and located within an active enhancer associated with the expression of Angiotensin II Receptor type 2 gene (*AGTR2*), a known modulator of insulin sensitivity. We further show that the risk T allele reduces binding of a nuclear protein, resulting in increased enhancer activity in muscle cells. Beyond providing novel insights into the genetics and pathophysiology of T2D, these results also underscore the value of reanalyzing publicly available data using novel analytical approaches.

---

During the last decade, hundreds of genome-wide association studies (GWAS) have been performed with the aim of providing a better understanding of the biology of complex diseases, improving their risk prediction, and ultimately discovering novel therapeutic targets^1^. However, the majority of the published GWAS just report the primary findings, which generally explain a small fraction of the estimated heritability. In order to better uncover the missing heritability, most strategies usually involve the generation of new genetic and clinical data. Very rarely, new studies are based on the revision and reanalysis of existing genetic data by applying more powerful analytic techniques and resources generated after the primary GWAS findings are published. These cost-effective reanalysis strategies are now possible, given the existence of (1) different data-sharing initiatives that gather large amounts of primary genotype and sequencing data for multiple human genetic diseases, as well as (2) new and improved GWAS methodologies and resources. Notably, genotype imputation with novel and denser sequence-based reference panels can now substantially increase the genetic resolution of GWASs from previously genotyped datasets^2^, reaching good quality imputation of low-frequency (minor allele frequency [MAF]: 0.01≤MAF<0.05) and rare variants (MAF<0.01). This increases the power to identify novel associations, as well as to refine known associated *loci* through fine-mapping. Moreover, the availability of public primary genetic data further allows the homogeneous integration of multiple datasets with different origins providing more accurate meta-analysis results, particularly at the low ranges of allele frequency. In contrast, common meta-analytic strategies usually involve the aggregation of summary statistics generated independently by different centers, each applying their own internal quality-control, data cleaning, genotype imputation, and association testing methodologies. Each of these steps represents a potential source of additional heterogeneity, which may ultimately result in loss of statistical power. Finally, the vast majority of reported GWAS analyses omits the X chromosome, even though it represents 5% of the genome and encodes for more than 1,500 genes^3,4^. The reanalysis of publicly available data also enables interrogation of this chromosome.

We hypothesized that a unified reanalysis of multiple publicly available datasets, applying homogeneous standardized QC, genotype imputation and association methods, as well as novel and denser sequence-based reference panels for imputation would provide new insights into the genetics and the pathophysiology of complex diseases. To test this we focused on type 2 diabetes (T2D), one of the most prevalent complex diseases for which many GWAS have been performed during the past decade^5,6,7,8,9,10,11^. These studies have allowed the identification of more than 100 independent *loci*, most of them driven by common variants, with a few exceptions^11,12,13^. Despite all these efforts, only a small fraction of the genetic heritability can be explained by established T2D *loci*, and the role of low-frequency variants, although recently proposed to be minor^14^, has still not been fully explored. The availability of large T2D genetic datasets in combination with larger and more comprehensive genetic variation reference panels^15,16,17,18^, gives us the opportunity to impute a significant increased fraction of low-frequency and rare variants, and to study their contribution to the risk of developing this disease. At the same time, this strategy also allows to fine map known associated *loci*, increasing the chances of finding causal variants and understanding their functional impact. We therefore gathered publicly available T2D GWAS cohorts with European ancestry, comprising a total of 13,857 T2D cases and 62,126 controls, to which we first applied harmonization and quality control protocols across the whole genome (including the X chromosome) and across samples, to later carry out imputation using 1,000 Genomes Project (1000G)^15^ and UK10K^16,17,18^ reference panels, as well as association testing. By using this strategy, we identified novel associated regions driven by common, low-frequency, and rare variants, fine-mapped and functionally annotated the existing and novel ones, and confirmed experimentally a regulatory mechanism disrupted by a novel rare and large effect variant identified at the X-chromosome.

## Results

### Overall analysis strategy

As shown in Figure 1, we first gathered all T2D case-control GWAS individual-level data that was available through the EGA and dbGaP databases (i.e. Gene Environment Association Studies [GENEVA], Welcome Trust Case Control Consortium [WTCCC], Finland-United States Investigation of NIDDM Genetics, Resource for Genetic Epidemiology Research on Aging [GERA] and Northwester NuGENE project [NuGENE]). We harmonized these cohorts, standardized quality control and the filtering of low-quality variants and samples (see Online Methods). After this process, a total of 70,127 subjects (70KforT2D; 12,931 cases and 57,196 controls; Supplementary Table 1) were retained for downstream analysis. Each of these cohorts was then imputed to the 1000G and UK10K reference panels using an integrative method, that selects for each variant the results from the reference panel that provides the highest accuracy, according to IMPUTE2 info score (see Online Methods). Finally, the results from each of these cohorts were meta-analyzed (Figure 1), obtaining a total of 15,115,281 variants with good imputation quality (IMPUTE2 info score ≥ 0.7, MAF ≥ 0.001, and I^2^ heterogeneity score < 0.75), across 12,931 T2D cases and 57,196 controls. Of these, 6,845,408 variants were common (MAF ≥ 0.05), 3,100,848 were low-frequency (0.01 ≤ MAF < 0.05) and 5,169,025 were rare (0.001 ≤ MAF < 0.01). Interestingly, the strategy of merging the imputation results derived from the UK10K and 1000G reference panels substantially improved the number of good quality imputed single nucleotide variants (SNVs) and insertion/deletions (*indels*), particularly within the low-frequency and rare spectrum, when compared with the imputation results obtained with each of the reference panels separately. For example, a set of 5,169,025 rare variants with good quality was obtained after integrating 1000G and UK10K results, while only 2,878,263 rare variants were imputed with 1000G and 4,066,210 with UK10K (Supplementary Figure 1 A). This strategy also allowed us to impute of 1,357,753 *indels* with good quality (Supplementary Figure 1 B).

**Figure 1:**
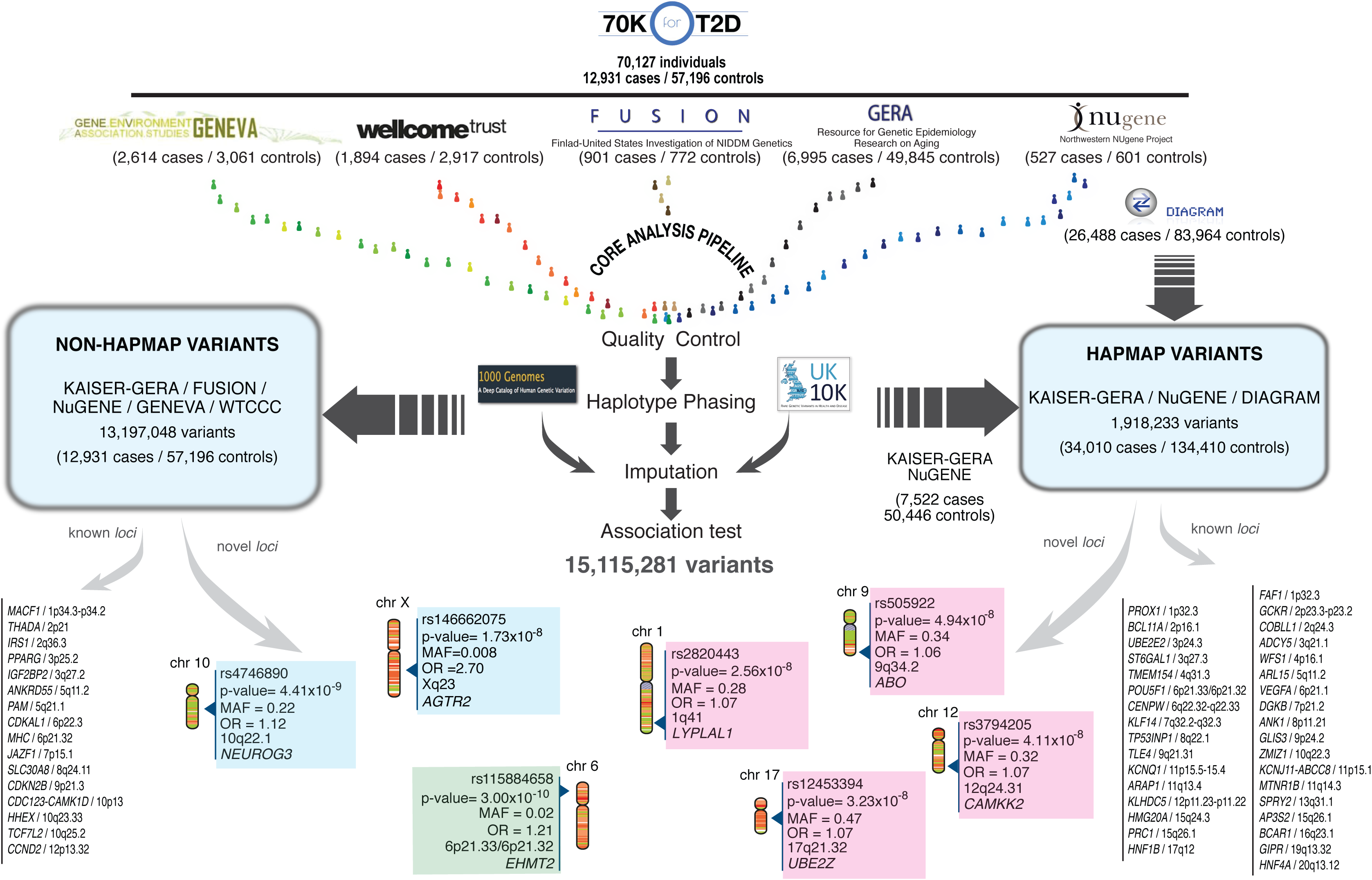
Discovery and replication strategy. Publicly available GWAS datasets representing a total of 12,931 cases and 57,196 controls (70KforT2D) were first quality controlled, phased and imputed using 1000G and UK10K separately. For those variants that were present in the DIAGRAM trans-ethnic meta-analysis we used the summary statistics to meta-analyze our results with the cohorts that had no overlap with any of the cohorts included in the DIAGRAM trans-ethnic meta-analysis. With this first meta-analysis we discovered four novel *loci* (within magenta panels). For the rest of the variants we meta-analyzed all the 70KforT2D datasets, which resulted in two novel *loci* (in blue panels). All the variants that were coding and that showed a p-value ≤ 1×10^−4^ were tested for replication by interrogating the summary statistics in the Type 2 Diabetes Knowledge Portal (T2D Portal) (http://www.type2diabetesgenetics.org/). This resulted in a novel low-frequency variant in the *EHMT2* gene (highlighted with a green panel).

In order to take full advantage of publicly available genetic data, we used three main meta-analytic approaches to adapt to three of the most common strategies for genetic data sharing: individual-level genotypes, summary statistics, as well as single-case queries through web services, like the Type 2 Diabetes Knowledge Portal (http://www.type2diabetesgenetics.org/). These three approaches allowed us to maximize the power of detection, by prioritizing largest sample sizes and best imputation quality, in particular for low-frequency, rare variants, and coding variants. We first meta-analyzed all summary statistics results from the DIAGRAM trans-ancestry meta-analysis^6^ (26,488 cases and 83,964 controls), selecting 1,918,233 common variants (MAF ≥ 0.05), mostly imputed from HapMap, with the corresponding fraction of non-overlapping samples in our 70KforT2D set, i.e. the GERA and the NuGENE cohorts, comprising a total of 7,522 cases and 50,446 controls (Figure 1, Supplementary Table 1). Second, the rest of variants (13,197,048), consisting of non-HapMap variants (mostly with MAF < 0.05) or not tested above, were meta-analyzed using all five cohorts that constitute the 70KforT2D resource (Supplementary Table 1). Finally, low-frequency variants located in coding regions and with p ≤ 1×10^−4^ were meta-analyzed using the non-overlapping fraction of samples with the data from the Type 2 Diabetes Knowledge Portal (http://www.type2diabetesgenetics.org) through the interrogation of exome array and whole-exome sequence data from ∼80,000 and ∼17,000 individuals, respectively^19,20,21^.

### T2D associated variants show enrichment of pathways involved in insulin response and pancreatic islet enhancers

As a first exploration of how our association results recapitulate the pathophysiology of T2D, we performed gene-set enrichment analysis with all the variants with p-value ≤ 1×10^−5^ using DEPICT^22^ (see Online Methods). This analysis showed enrichment of genes expressed in pancreas (ranked 1^st^ in tissue enrichment analysis, *p*=7.8×10^−4^, FDR<0.05, Supplementary Table 2) and cellular response to insulin stimulus (ranked 2^nd^ in gene set enrichment analysis, *p*=3.9×10^−8^, FDR=0.05, Supplementary Table 3, Supplementary Figure 2, Supplementary Figure 3), in concordance with the current knowledge of the molecular basis of T2D.

In addition, variant set enrichment analysis of the T2D-associated credible sets across regulatory elements defined in isolated human pancreatic islets showed a significant enrichment for active regulatory enhancers (Supplementary Figure 4), suggesting that causal SNPs within associated region have a regulatory function, as previously reported^23^.

### Identification, fine-mapping and functional characterization of novel and previously known loci

The three association strategies allowed us to finally identify 57 genome-wide significant associated *loci* (*p*≤5×10^−8^), of which seven were not previously reported as associated with T2D (Table 1). The remaining 50 *loci* were already known, and included, for example, two low-frequency variants recently discovered in Europeans, one located within one of the *CCND2* introns (rs76895963) and a missense variant within the *PAM*^13^ gene. Furthermore, as a quality control of these results, we confirmed that the magnitude and direction of the effect of all the associated variants (*p*≤0.001) were highly consistent with those reported previously (Rho=0.92, *p*=1×10^−248^, Supplementary Figure 5). In fact, the direction of effect was consistent with all 139 previously reported variants except three that were discovered in east and south Asian populations (Supplementary Table 4).

**Table 1.**
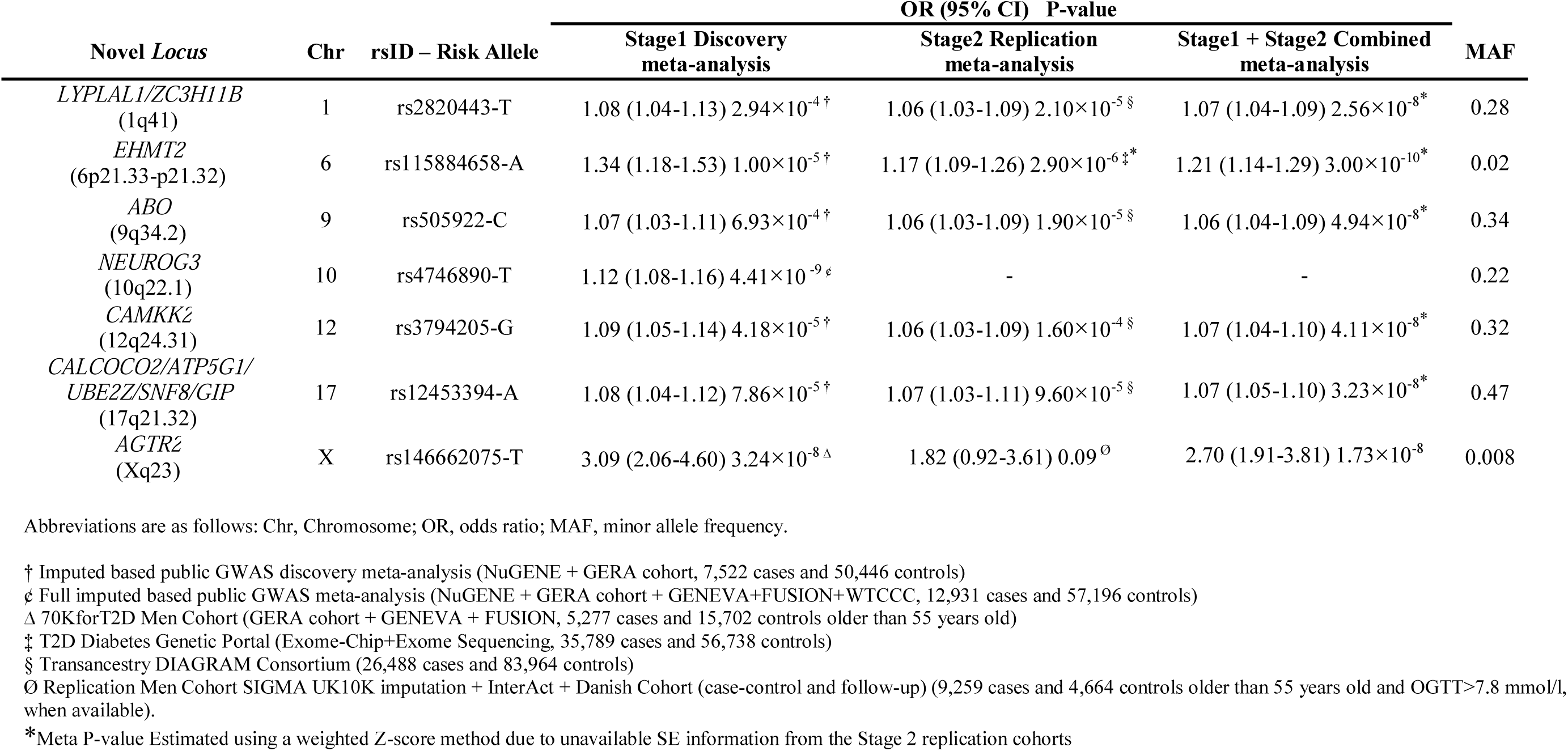
Novel T2D associated *loci*.

The high coverage of genetic variation achieved in this study allowed us to fine-map known and novel *loci*, providing more candidate causal variants for downstream functional interpretations. For this, we constructed 99% credible variant sets^24^, -i. e. the subset of variants that have, in aggregate, 99% probability of containing the true causal variant-for all 57 *loci* (Supplementary Table 5). As an important improvement over previous T2D genetic studies, we identified small structural variants within the credible sets, consisting mostly of insertions and deletions between 1 and 1,975 nucleotides. In fact, out of the 8,348 variants included within the credible sets for these *loci*, 927 (11,1%) were *indels*, of which 105 were genome-wide significant (Supplementary Table 6). Interestingly, by integrating imputed results from 1000G and UK10K reference panels we gain up to 41% of *indels*, which were only identified by one of the two reference panels, confirming the advantage of integrating the results from both reference panels. Interestingly, 15 of the 71 previously reported *loci* we replicated (*p*≤ 5.3×10^−4^ after correcting for multiple testing), have an *indel* as the top variant, highlighting the potential role of this type of variation in the susceptibility for T2D. For example, within the *IGF2BP2* intron, a well-established and functionally validated *locus* for T2D^25,26^, we found that 12 of the 57 variants within its 99% credible set correspond to *indels* with genome-wide significance (5.6×10-^16^ < *p* < 2.4×10^−15^), which collectively represent 18.4% posterior probability of being causal.

To prioritize causal variants within all the identified associated *loci*, we annotated their corresponding credible sets using the variant effector predictor (VEP) for coding variants^27^ (Supplementary Table 7), and the Combined Annotation Dependent Depletion (CADD) tool for non-coding variation (Supplementary Table 8)^28^. Additionally, we tested the effect of all variants on expression across multiple tissues by interrogating GTEx^29^ and RNA-sequencing gene expression data from pancreatic islets^30^.

### Identification of new signals driven by common variants

Beyond the detailed characterization of the known T2D associated regions, we also identified novel *loci*, among which, five were driven by common variants linked to modest effect sizes (1.06 < OR <1.12; Table 1, Figure 2, Supplementary Figure 6 and 7).

**Figure 2:**
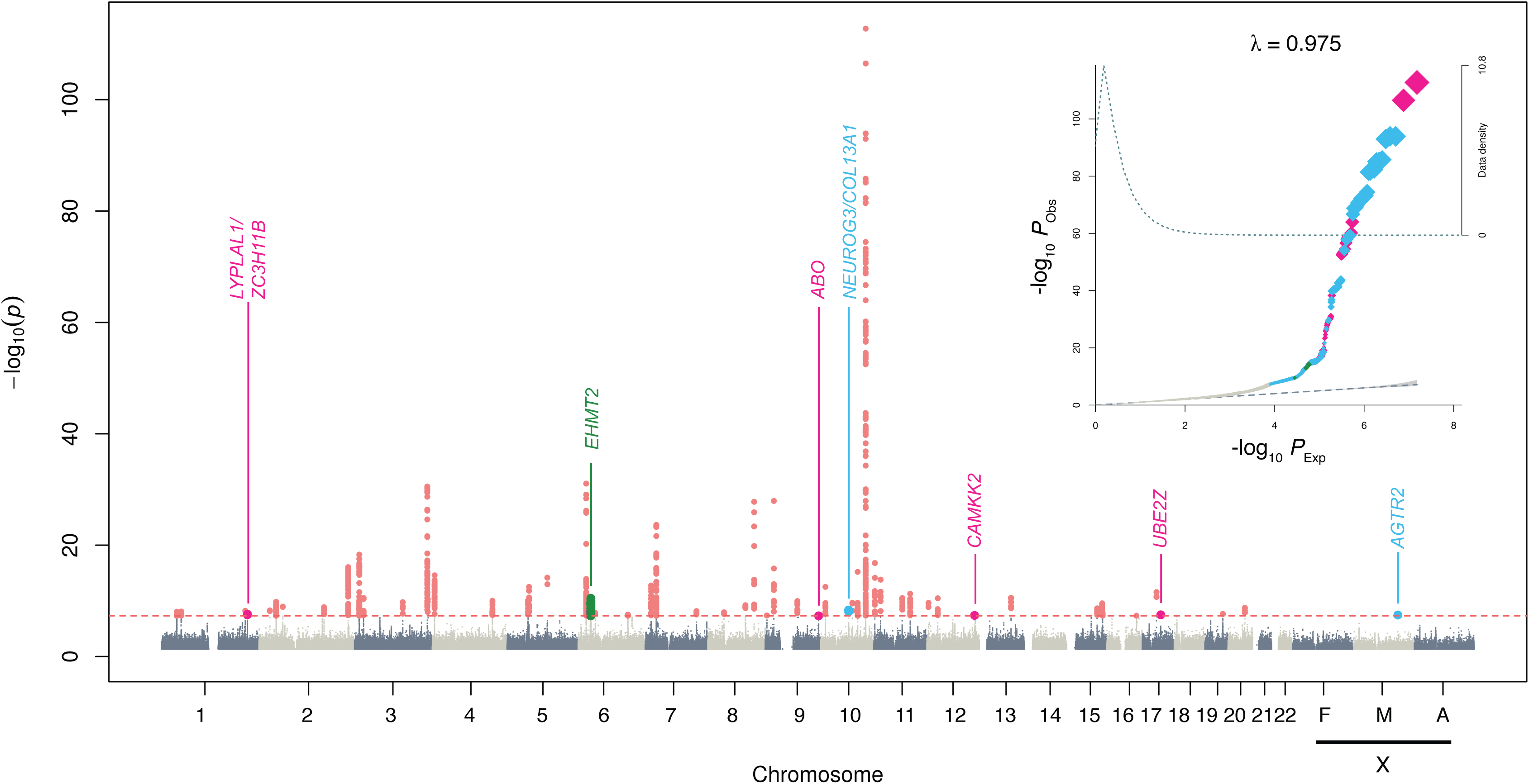
Manhattan and Quantile-Quantile plot (QQ-plot) of the discovery and replication genome-wide meta-analysis. The upper corner represents the QQ-plot. Expected −log10 p-values under the null hypothesis are represented in the x-axis while observed −log_10_ p-values are represented in the y-axis. Observed p-values were obtained according to the suitable replication dataset used (as shown in Figure 1) and were depicted using different colors. HapMap variants were meta-analyzed using the Trans-Ethnic summary statistics from the DIAGRAM study and our meta-analysis based on the Genetic Epidemiology Research on Aging (GERA) cohort and the Northwestern NuGENE Project and that resulted in novel associations depicted in magenta. The rest of non-HapMap variants meta-analyzed using the full 70KforT2D cohort are represented in grey, highlighting in light blue the fraction of novel GWAS-significant variants. Coding low-frequency variants meta-analyzed using the 70KforT2D and the T2D Portal data that resulted in novel GWAS-significant associations are depicted in green. The shaded area of the QQ-plot indicates the 95% confidence interval under the null and a density function of the distribution of the p-values was plotted using a dashed line. The λ is a measure of the genomic inflation and corresponds to the observed median χ^2^ test statistic divided by the median expected χ^2^ test statistic under the null hypothesis. The Manhattan plot, representing the −log10 p-values were colored as explained in the QQ-plot. All known GWAS-significant associated variants within known T2D genes are also depicted in red. X-chromosome results for females (F), males (M) and all individuals (A) are also included.

Within the first novel T2D-associated *locus* in chromosome 1q41 (*LYPLAL1-ZC3H11B*, rs2820443, OR=1.07 [1.04-1.09], *p*=2.6×10^−8^), several variants have been previously associated with waist-to-hip ratio in females, height, visceral adipose fat in females, adiponectin levels, fasting insulin, and non-alcoholic fatty liver disease^31,32,33,34,35,36^. Among the genes of this *locus, LYPLAL1*, which encodes for lysophospholypase-like 1, appears as the most likely effector gene, as it has been found to be downregulated in mouse models of diet-induced obesity and upregulated during adipogenesis^37^.

Second, a novel *locus* at chromosome 9q34.2 region (*ABO*, rs505922, OR=1.06 [1.04-1.09], *p*=4.9×10^−8^) includes several variants that have been previously associated with other metabolic traits. For example, the variant rs651007, in linkage disequilibrium (LD) with rs505922 (r^2^=0.507), has been recently associated with fasting glucose^38^; and rs514659 (r^2^ with top=1) is associated with an increased risk for cardio-metabolic disorders^39^. Interestingly one of the variants within the credible set is the one base-pair frame-shift deletion defining the blood group O^40^. In concordance with previous results that linked O blood type with a lower risk of developing T2D^41^, the frameshift deletion determining the blood group type O was associated with a protective effect for T2D in our study (rs8176719, *p*=3.4×10^−4^, OR=0.95 [0.91-0.98]). In addition, several variants within this credible set are associated with the expression of the *ABO* gene in multiple tissues including skeletal muscle and adipose tissue, and pancreatic islets (Supplementary Table 9, Supplementary Table 10).

Third, a novel *locus* at chromosome 10q22.1 *locus* (*NEUROG3/COL13A1/RPL5P26*, rs2642587, OR=1.12 [1.08-1.16], *p*=8.4×10^−9^), includes *NEUROG3* (Neurogenin3), which is an essential regulator of pancreatic endocrine cell differentiation^42,43^. Mutations in this gene have thus been reported to cause permanent neonatal diabetes^44^, but a role of this gene in T2D has not been previously reported^45^.

The lead common variant of the fourth novel *locus* at chromosome 12q24.31 (rs3794205, OR=1.07 [1.04-1.10], *p*=4.1×10^−8^), lies within an intron of the *CAMKK2* gene, previously implicated in cytokine-induced beta cell death^46^. However other variants within the corresponding credible set could be, in fact, responsible for the molecular action behind this association, like a missense variant within the *P2RX7*, a gene previously associated with glucose homeostasis in humans and mice^47^; or other variant (rs11065504, r^2^ with lead variant=0.81) that is associated with the regulation of the *P2RX4* gene in tibial artery and in whole blood according to GTEx (Supplementary Table 9).

The fifth novel *locus* driven by common variants is located within 17q21.32 (rs12453394, OR=1.07 [1.05-1.10], *p*=3.23×10^−8^). It involves three missense variants located within the *CALCOCO2, SNF8* and *GIP* genes. *GIP* encodes for glucose-dependent insulinotropic peptide, a hormonal mediator of enteral regulation of insulin secretion^48^. Variants in the GIP receptor (GIPR) have previously been associated with insulin response to oral glucose challenge and beta-cell function^49^. *GIP* is thus a plausible candidate effector gene of this *locus*^50^.

### Identification of a new signal driven by a low-frequency variant

We selected all low-frequency (0.01≤MAF<0.05) variants with *p*≤ 1×10^−4^ in the 70KforT2D meta-analysis that were annotated as altering protein-coding variants according to VEP. This resulted in 15 coding variants that were meta-analyzed with exome array and whole-exome sequencing data from a total of ∼97,000 individuals^19, 20, 21^ after excluding the overlapping cohorts between the different datasets. This analysis highlighted a novel genome-wide association driven by a low-frequency missense variant (Ser58Phe) within the *EHMT2* gene at chromosome 6p21.33 (rs115884658, OR=1.21 [1.14-1.29], *p*=3.00×10^−10^; Figure 2, Supplementary Figures 6 and 7). *EHMT2* is involved in the mediation of FOXO1 translocation induced by insulin^51^. Since this variant was less than 1 Mb away from *HLA-DQA1*, which was recently reported to be associated with T2D^52^, we performed a series of reciprocal conditional analyses and excluded the possibility that our analysis was capturing previously reported T2D^52^ or T1D^10,53,54^ signals (Supplementary Table 11). Beyond this missense *EHMT2* variant, other low-frequency variants within the corresponding credible set may also be causal. For example, rs115333512 (r^2^ with lead variant=0.28) is associated with the expression of *CLIC1* in several tissues according to GTEx (multi-tissue Meta-Analysis *p*=8.9×10^−16^, Supplementary Table 9). In addition, this same variant is also associated with the expression of the first and second exon of the *CLIC1* mRNA in pancreatic islet donors^30^ (*p*(exon 1)=1.4×10^−19^; *p*(exon 2)=1.9×10^−13^; Supplementary Table 10). Interestingly, *CLIC1* has been reported as a direct target of metformin by mediating the anti-proliferative effect of this drug in human glioblastoma^55^. All these findings support *CLIC1*, as an additional possible effector transcript, likely driven by rs115333512.

### Identification of a novel rare variant in the X chromosome associated with 2.7-fold increased risk for T2D

As for many other complex diseases, the majority of published large-scale T2D GWAS studies have omitted the analysis of the X chromosome, with the notable exception of the identification of a T2D associated region near the *DUSP9* gene in 2010^56^. To fill this gap, we tested the X chromosome genetic variation for association with T2D. To account for heterogeneity of the effects and for the differences in imputation performance between males and females, the association was stratified by sex and tested separately, as well as together by meta-analyzing the results from males and females. This analysis was able to replicate the *DUSP9 locus*, not only through the known rs5945326 variant (OR=1.15, *p*=0.049), but also through a three-nucleotide deletion located within a region with several promoter marks in the liver (rs61503151 [G/GCCA], OR=1.25, *p*=3.5×10^−4^), and in high LD with the first reported variant (R^2^=0.62). Conditional analyses showed that the originally reported variant was no longer significant (OR=1.01, *p*=0.94) when conditioning by the newly identified variant, rs61503151. On the other hand, when conditioning on the previously reported variant, rs5945326, the effect of the newly identified *indel* remained significant and with higher effect size (OR=1.33, *p*=0.003), placing this deletion, as a more likely candidate causal variant for this *locus*.

In addition, we identified a novel genome-wide significant signal in males at the Xq23 *locus* driven by a rare variant (rs146662075, MAF=0.008, OR=2.94 [2.00-4.31], *p*=3.5×10^−8^; Figure 3A). We tested the accuracy of the imputation of this variant by comparing the imputed results from the same individuals genotyped by two different platforms (see Online Methods) and found that the imputation for this variant was highly accurate in males, and when using UK10K, but not in females, or when using 1000G (R^2^ [UK10K,males]=0.94, R^2^ [UK10K,females]=0.66, R^2^ [1000G,males]=0.62, R^2^ [1000G,females]= 0.43, Supplementary Figure 8). Whether this association is specific to men or whether it also affects female carriers remains to be clarified, since the imputation for this variant in females was not accurate enough.

**Figure 3:**
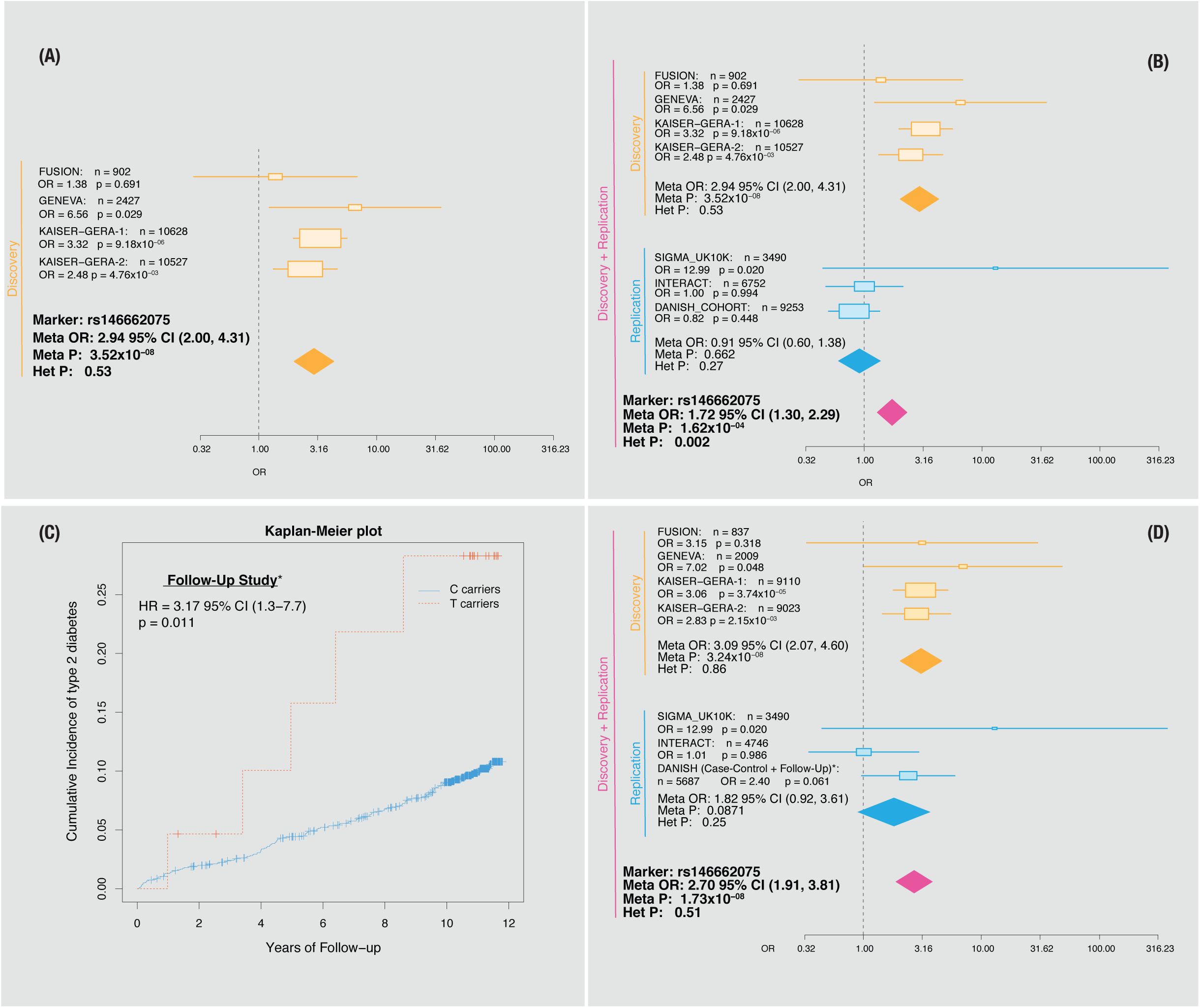
Discovery and replication of rs14666075 association signal. Forest plot for the rs146662075 variant using (A) the discovery cohorts and when including also (B) the replication datasets. Cohort-specific odds ratios are denoted by boxes proportional to the size of the cohort and 95% CI error bars. The combined OR estimate for all the datasets is represented by a green diamond, where the diamond width corresponds to 95% CI bounds. The p-value for the meta-analysis (Meta P) and for the heterogeneity (Het P) of odds ratio is shown. C) Plot showing the cumulative incidence of type 2 diabetes for an 11 years follow-up. The red line represents the T carriers and in light blue, C carriers are represented (n=1,652, cases=158.). D) Forest plot after excluding controls younger than 55 years old and OGTT>7.8 mmol/l in both the discovery and replication cohorts when available.

To further validate this association and to discard potential imputation artifacts, we next analyzed two independent cohorts by performing imputation with the UK10K reference panel (SIGMA^57^, INTERACT^58^), and a third cohort by *de-novo* genotyping the rs146662075 variant in several Danish sample sets. The initial meta-analysis, including the three replication datasets did not reach genome-wide significance (OR=1.72, *p*=1.6×10^−4^) (Figure 3B), and revealed a strong degree of heterogeneity (heterogeneity *p*_het_=0.002), which appeared to be driven by the replication cohorts. Within one of the case-control studies there was a nested cohort study; this study, the Inter99 prospective cohort, consisted of 1,652 non-diabetic male subjects, of whom 158 developed T2D after eleven years of follow-up. Analysis of incident diabetes in this cohort, confirmed the association with the same allele as previously seen in the case-control studies, with carriers of the rare T allele having increased risk of developing incident diabetes, compared to the C carriers (Cox-proportional Hazards Ratio (HR)=3.17 [1.3-7.7], *p*=0.011, Figure 3C): in fact, nearly 30% carriers of the T risk allele developed incident T2D during 11 years of follow-up, compared to only ∼10% of noncarriers.

In order to gain further support for this association, we thoroughly compared the clinical and demographic characteristics of the discovery and replication datasets in an attempt to understand the strong degree of heterogeneity observed when adding the replication datasets. We found that some of the replication datasets contained control subjects that were significantly younger than the average age at onset of T2D reported in this study and in Caucasian populations in general^59^. This was particularly clear for the Danish cohort (age controls [95%CI] = 46.9 [46.6-47.2] vs age cases [95%CI] = 60.7 [60.4-61.0]) and for Interact (age controls [95%CI] = 51.7 [51.4-52.1] vs age cases [95%CI] = 54.8 [54.6-55.1]) (Supplementary Figure 9). Since in the Inter99 prospective cohort we observed that 30% of subjects developed incident T2D over 11 years of follow-up, we performed an additional analysis using a stricter definition of controls, in order to avoid the presence of pre-diabetes or individuals that may further develop diabetes after reaching the average age at onset. We therefore applied two additional exclusion criteria in order to obtain a stricter set of controls: (i) excluding subjects younger than 55 years old, and (ii) restricting the analysis, when possible, to individuals with measured 2 hours plasma glucose values during oral glucose tolerance test (OGTT) below 7.8 mmol/l, which is employed to diagnose impaired glucose tolerance (pre-diabetes), a strong risk factor for developing T2D^60^. We repeated the metaanalysis using the first filtering criterion for controls for both the discovery and replication datasets, including only controls older than 55. While the application of the first filter did not yet provide genome-wide significant results (Supplementary Figure 10), when we added the second filter (only possible in the Danish cohort), we obtained replication results consistent with the initial discovery results. Moreover, we also integrated the Cox-proportional hazards results into the meta-analysis by using a method that accounts for overlapping subjects (MAOS)^61^ allowing us to integrate the longitudinal (follow-up) and the case-control study. This final meta-analysis confirmed the association at rs146662075 resulting in genome-wide significance and without significant heterogeneity (OR=2.7 (1.91, 3.81), *p*=1.7×10^−08^, phet=0.51, Figure 3D). These results therefore support the existence of a genetic association with T2D is driven by a rare variant at X chromosome.

### The rs146662075 T risk allele is associated with 5-fold greater enhancer activity and disruption of allele specific nuclear protein binding

We next explored the possible molecular mechanism behind this association by using different genomic resources and experimental approaches. The credible set of this region contained three variants, with the leading SNP alone (rs146662075), showing 78% posterior probability of being causal (Supplementary Figure 7, Supplementary Table 5), as well as the highest CADD score (scaled C-score=15.68, Supplementary Table 8). rs146662075 lies within a chromosomal region enriched in regulatory (DNAse I) and active enhancer (H3K27ac) marks, between the *AGTR2* (at 103 kb) and the *SLC6A14* (at 150Kb) genes. Interestingly, the closest gene, *AGTR2*, which encodes for the angiotensin II receptor type 2, has been previously associated with insulin secretion and resistance^62,63,64^. Analysis of publicly available epigenomic datasets^65^ showed that this region lacks H3K27ac enrichment in human islet chromatin, whereas there is a positive correlation between the H3K27ac enhancer marks and the expression of *AGTR2* across multiple tissues, showing the highest signal of both H3K27ac and *AGTR2* RNA-seq (but not *SLC6A14*) expression in fetal muscle (Figure 4A).

**Figure 4:**
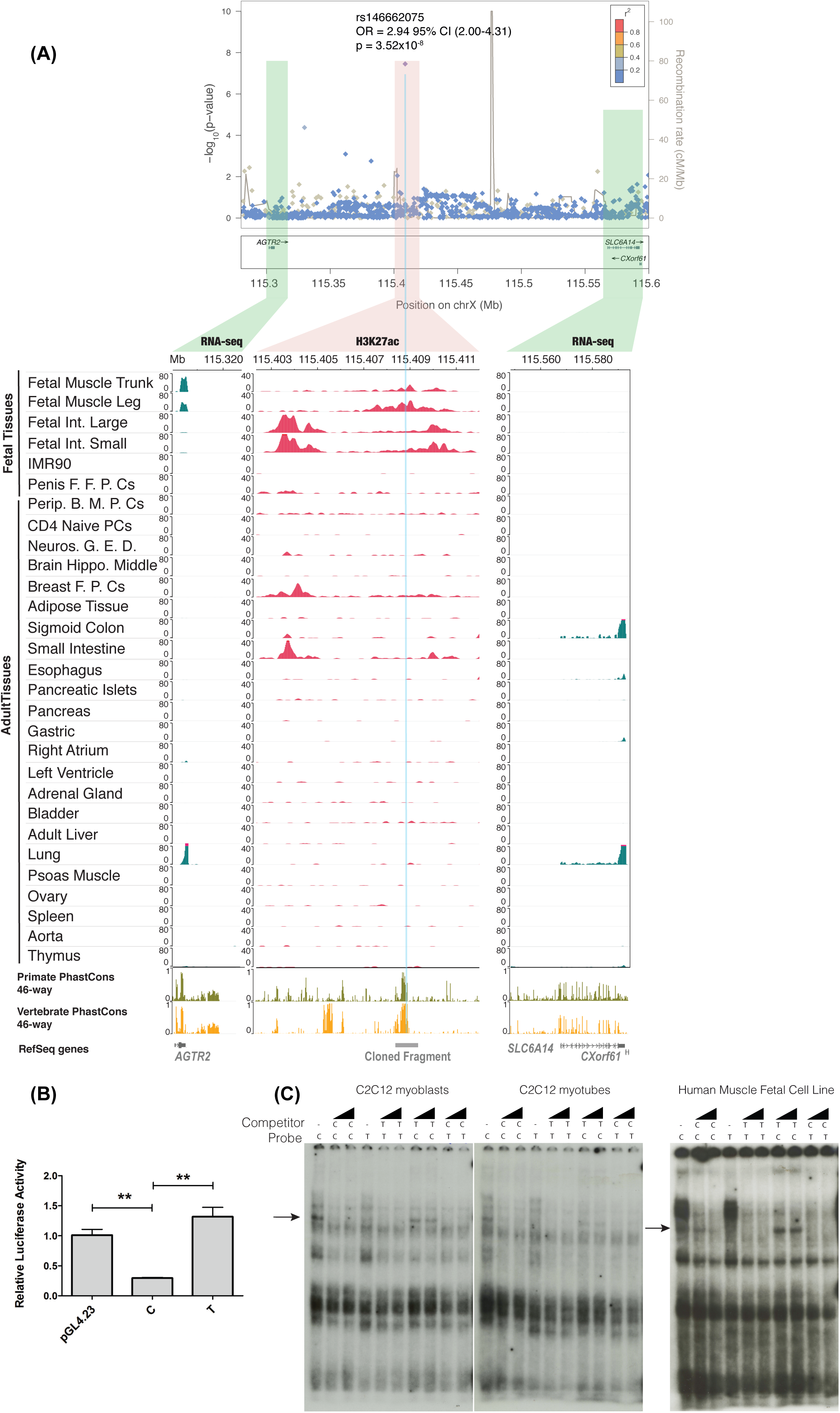
Functional characterization of rs146662075 association signal. A) Signal plot for X chromosome region surrounding rs146662075. Each point represents a variant, with its *p*-value (on a −log10 scale, y-axis) derived from the meta-analysis results from association testing in males. The x-axis represents the genomic position (Hg19). Below, representation of H3K27ac and RNA-seq in a subset of cell-types for which RNA-seq and H3K27ac was available is also shown. The association between RNA-seq signals and H3K27ac marks suggest that *AGTR2* is the most likely regulated gene by the enhancer that harbors rs146662075. B) The presence of the common allelic variant rs146662075-C reduces enhancer activity in luciferase assays performed in a mouse myoblast cell line. C) Electrophoretic mobility shift assay in C2C12 myoblast cell lines, C2C12 differentiated myotubes and human fetal myoblasts showed allele-specific binding of a ubiquitous nuclear complex. The arrows indicate the allele-specific binding event. Competition was carried out using 50- and 100-fold excess of corresponding unlabeled probe.

We next studied whether the region encompassing the rs146662075 variant could act as a transcriptional enhancer and whether its activity was allele-specific. For this, we linked the DNA region, with either the T (risk) or the C (common) allele, to a minimal promoter and performed luciferase assays in a mouse myoblast cell line. The luciferase analysis showed an average 4.4 fold increased activity for the disease-associated T allele, compared to the expression measured with the common C allele, suggesting an activating function of the T allele, or a repressive function of the C allele (Figure 4B). Consistent with these findings, electrophoretic mobility shift assays using nuclear protein extracts from mouse myoblast cell lines, differentiated myotubes, and human fetal muscle cell line, revealed sequence-specific binding activity of the C allele, but not of the rare T allele (Figure 4C). Overall, these data indicate that the risk T allele prevents binding by a nuclear protein that is associated with decreased activity of an *AGTR2*-linked enhancer.

## Discussion

To date, a large fraction of GWAS datasets have been made publicly available through repositories such as dbGaP and EGA. However, how these resources are useful to increase the knowledge of the genetic factors contributing to disease susceptibility has not been fully explored. In this study, we demonstrate the utility of these publicly available resources. By harmonizing and reanalyzing existing and publicly available T2D GWAS data, and by performing genotype imputation with two whole-genome sequence-based reference panels, we have been able to explore deeply the genetic architecture of T2D. This strategy allowed us to impute and test for association with T2D more than 15 million of high quality imputed variants, including low-frequency, rare, and small insertions and deletions across chromosomes 1 to 22 and X.

The reanalysis of these data confirmed a large fraction of already known T2D *loci*, for which we identify and propose novel potential causal variants by fine-mapping and functionally annotating each *locus*. Our in-depth characterization of each of these *loci* reached an improved accuracy over prior efforts, providing for the first time a comprehensive coverage of structural variants, which point to previously unobserved candidate causal variants in known and novel *loci*.

This reanalysis also allowed us to identify seven novel associations driven by common variants in or near *LYPLAL1, NEUROG3, CAMKK2, ABO* and *GIP;* a low-frequency variant in *EHMT2*, and one rare variant in the X chromosome. This rare variant identified in Xq23 chromosome, was located near the *AGTR2* gene, and showed a nearly three-fold increased risk for T2D in males, which represents, to our knowledge, the largest effect size identified so far in Europeans, and complements other large effect size variants identified in other specific populations^11,12^.

This study also highlights the importance of a strict classification of both cases and controls in order to identify rare variants associated with disease, as our initial discovery for the Xq23 *locus* was not replicated when using the standard classification of cases and controls (based on fasting plasma glucose or glycated hemoglobin, HbA1c), but only when reclassifying the control group by restricting it to non-diabetic individuals older than 55 years (average age at onset of T2D), and with confirmed normal glucose tolerance using the stricter OGTT measure. This is in line with previous results obtained for a T2D population specific variant found in Inuit within the *TBC1D4* gene, which was only significant when using OGTT as criteria for classifying cases and controls, but not when using HbA1c^11^. The use of this strict criterion for T2D diagnosis was in fact very clinically relevant, as 32% carriers of the *TB1CD4* variant would remain undiagnosed T2D or prediabetes patients when OGTT was not assessed^66^. Despite the need for a stricter definition of controls in our replication case-control study, our results were further confirmed by the observation that 30% of the rs146662075 risk allele carriers developed T2D over 11 years of follow-up, compared to 10% of non-carriers, and suggest that early identification of these subjects by genotyping this variant may be useful to tailor pharmacological or lifestyle intervention in order to prevent or delay their onset of T2D.

Using binding and gene-reporter analyses we demonstrate a functional role of this variant and propose a possible mechanism behind the pathophysiology of T2D in T risk allele carriers, where this rare variant could favor a gain of function of *AGTR2*, which has been previously associated with insulin resistance^62^. *AGTR2* appears, therefore, as a potential therapeutic target for this disease, which would be in line with previous studies that showed that the blockade of the renin-angiotensin system in mice^67^ and in humans^68^ prevents the onset of type 2 diabetes and restores normoglycemia^69,70^.

Overall, beyond our significant contribution to the understanding of the molecular basis of T2D, our study also highlights the potential of the reanalysis of public data, as a complement to large studies using newly-generated data. This study will inform the open debate in favor of pushing data-sharing and democratization initiatives^71,72^, highlighting their importance for the study of the genetics and pathophysiology of complex diseases, which may lead to new preventive and therapeutic applications.

## Methods

Online Content Methods, along with any additional Extended Data display items, are available in the online version of the paper; references unique to these sections appear only in the online paper.

## Data availability

The complete summary statistics are deposited at the Type 2 Diabetes Knowledge portal (www.type2diabetesgenetics.org/).70KforT2D: Identification of novel T2D *loci* with publicly available GWAS data

## Acknowledgements

This work has been sponsored by the grant SEV-2011-00067 of Severo Ochoa Program, awarded by the Spanish Government. This work was supported by an EFSD/Lilly research fellowship. Josep M. Mercader was supported by Sara Borrell Fellowship from the Instituto Carlos III. Sílvia Bonàs was FI-DGR Fellowship from FI-DGR 2013 from Agència de Gestió d’Ajuts Universitaris i de Recerca (AGAUR, Generalitat de Catalunya). This study makes use of data generated by the Wellcome Trust Case Control Consortium. A full list of the investigators who contributed to the generation of the data is available from www.wtccc.org.uk. Funding for the project was provided by the Wellcome Trust under award 076113. This study also makes use of data generated by the UK10K Consortium, derived from samples from UK10K COHORT IMPUTATION (EGAS00001000713). A full list of the investigators who contributed to the generation of the data is available in www.UK10K.org. Funding for UK10K was provided by the Wellcome Trust under award WT091310. We acknowledge PRACE for awarding us access to MareNostrum supercomputer, based in Spain at Barcelona. The technical support group from the Barcelona Supercomputing Center is gratefully acknowledged. This project has received funding from the European Union’s Horizon 2020 research and innovation programme under grant agreement No 667191. Mercè Planas-Fèlix is funded by the Obra Social Fundación la Caixa fellowship under the Severo Ochoa 2013 program. Irene Miguel-Escalada has received funding from the European Union's Horizon 2020 research and innovation programme under the Marie Sklodowska-Curie grant agreement No 658145. The Novo Nordisk Foundation Center for Basic Metabolic Research is an independent Research Center at the University of Copenhagen partially funded by an unrestricted donation from the Novo Nordisk Foundation (www.metabol.ku.dk). We also thank Marcin von Grotthuss for their support for uploading the summary statistics data to the Type 2 Diabetes Genetic Portal (AMP-T2D portal). Finally,we thank all the Computational Genomics group at the BSC for their helpful discussions and valuable comments on the manuscript.

